## Supporting Information for "Differential Crosslinking and Contractile Motors Drive Nuclear Chromatin Compaction"

### 1. COMPUTATIONAL MODEL

The chromatin-lamina system is modeled with chromatin represented as a Rouse polymer chain and the nuclear lamina as an elastic, polymeric spherical shell. Linkages are formed between the chromatin and the lamina. The shell consists of 5000 monomers positioned using a Fibonacci sphere algorithm, with each pair of neighboring monomers connected by Hookean springs of spring constant $K$, forming a mesh with an average coordination number of 4.5. The chromatin is modeled as a Rouse chain with 5000 monomers ($N$), generated via a three-dimensional self-avoiding random walk on an FCC lattice. Each monomer has a radius $r_p = 0.43089$ and experiences soft-core repulsion to capture excluded volume effects. The lamina monomers share the same physical properties (size and spring strength) as the chromatin monomers. To initialize the system, the chromatin is confined within a spherical shell of radius R$s = 10$. The shell is gradually shrunk by moving lamina monomers inward, which compresses the chromatin via steric interactions. Once the desired radius is reached, spring lengths and monomer positions are adjusted to finalize the initial configuration (Fig. S1). This process is repeated to generate 50 independent initial configurations. Given $r_p = 0.43089$, the packing fraction in the confined shell is approximately $\phi_{\text{pack}} \approx 0.4$ (details of simulation parameters are provided in Table S1).

Chromatin crosslinks are introduced by adding $N_c$ springs between randomly selected pairs of chromatin monomers, provided their mutual distance is less than $r_{\text{link}} = 3r_p$. The stiffness of the crosslink springs matches that of the polymer backbone. Motor activity is incorporated by designating $N_m$ chromatin monomers as active. Each active monomer exerts a force $\boldsymbol{F_a} = \pm f_m \hat{r}_{ij}$ on nearby monomers, where $f_m$ denotes the motor force magnitude, and $\hat{r}_{ij}$ is the unit vector between the interacting monomers. We consider two types of motors: extensile, which push monomers apart, and contractile, which draw them together.

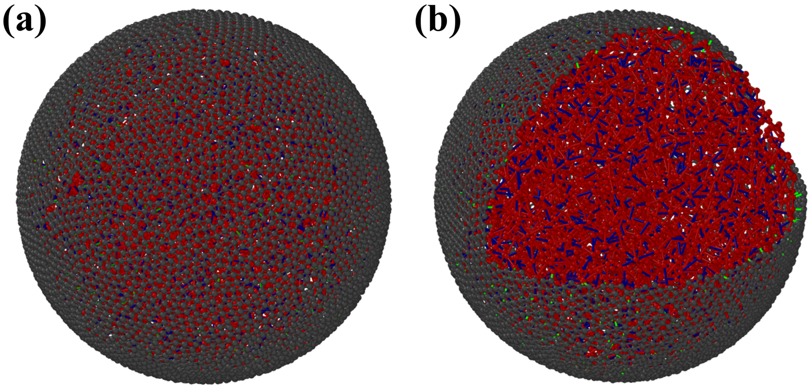


Fig. S1: *Snapshots of the initial configuration of the chromatin–lamina system.* (a) Initial configuration of chromatin confined within the nuclear lamina. (b) Crosslinked chromatin is shown with red beads; crosslinks are depicted in blue, and chromatin-lamina linkages are highlighted in green.

*
†
‡

| Parameters | Numerical Value |
|---|---|
| Number of polymer monomers ($N$) | 5000 |
| Number of shell particles ($N_s$) | 5000 |
| Radius of a polymer monomer ($r_p$) | 0.43089 |
| Radius of a shell particle ($r_s$) | 0.43089 |
| Radius of the hard shell ($R_s$) | 10 |
| Packing fraction ($\phi_{pack}$) | 0.4 |
| Number of crosslinks ($N_c$) | 0, 500, 1000, 2000 |
| Number of linkages ($N_L$) | 0, 50, 250, 600 |
| Number of motors ($N_m$) | 500 |
| Magnitude of motor force ($f_m$ ) | 10 |
| Motor turnover time scale ($\tau_m$) | 20 |
| Harmonic spring constant ($K$) | 140 |
| Excluded volume strength ($K_{Ex}$) | 140 |
| Damping ($\xi$) | 1 |
| Simulation timestep ($d\tau$) | $10^{-4}$ |
| Thermal Energy ($K_BT$) | 1 |
| Diffusion constant ($D$) | 1 |
| Cutoff used for crosslinks ($r_{\text{link}}$) | $3\times r_p$ |
| Motor radius ($r_{\text{motor}}$) | $1.5\times r_p$ |

**Table S1:** Parameters used in the simulations.

### 2. SIMULATION RESULTS

#### 2.1. Radius of gyration of chromatin polymer and radial density distribution of chromatin monomers

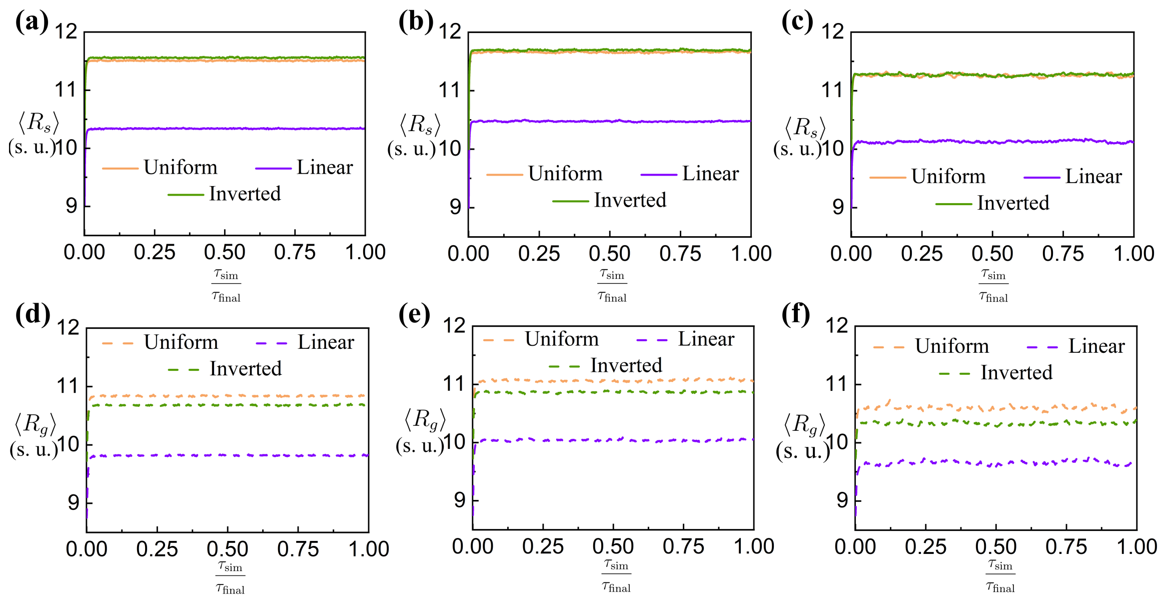


Fig. S2: *Temporal profile of nuclear radius and chromatin radius of gyration for passive, extensile, and contractile motors across different crosslink profiles.* Time evolution of the average radius of the nuclear lamina, $\langle R_s \rangle$ (solid lines, top row), and the chromatin radius of gyration, $\langle R_g \rangle$ (dashed lines, bottom row), for (a) passive, (b) extensile, and (c) contractile motor cases with different crosslink density profiles. The results shown correspond to $N_c = 2000$ crosslinks and $N_L = 600$ chromatin-lamina linkages. All quantities are reported in simulation units (s. u.), where 1 s. u. of length = 1 $\mu m$ and 1 s. u. of time = 0.5 s.

For a chromatin polymer, the radius of gyration is defined as $R_g = \sqrt{\frac{1}{N}\sum_{i=1}^{N}(\boldsymbol{r}_i - \boldsymbol{r}_{\mathrm{com}})^2}$, where $N = 5000$ is the total number of monomers in the chain, $r_i$ is the position of $i^{th}$ monomer, and $r_{\mathrm{com}}$ is the center of mass of the chromatin chain. The radius of the nuclear shell is fixed at $R_s = 10$ considering as a initial hard shell. The soft shell expands in response to thermal fluctuations and the active forces generated by the chromatin chain confined within the nucleus. Fig. S2 shows the average nuclear radius, $\langle R_s \rangle$ (top row, solid lines), and average radius of gyration, $\langle R_g \rangle$, of the chromatin chain (bottom row, dashed lines) as a function of simulation time. Following a short initial expansion, the radii of both the chromatin chain and the nuclear shell reach a plateau by $100\tau$, indicating the onset of a steady-state configuration.

#### 2.2. Influence of chromatin-lamina linkages on chromatin spatial distribution

We consider a reference case in which both chromatin crosslinks ($N_c$) and chromatin–lamina linkages ($N_L$) are absent. The resulting radial chromatin distributions for passive, extensile, and contractile motor cases are shown in Fig. S3a. In the absence of crosslinks and lamina linkages, the chromatin density remains largely uniform across the nuclear radius, $R_s$. A slight accumulation of chromatin is observed toward the nuclear interior, which becomes more pronounced in the presence of contractile motor activity (Fig. S3a).

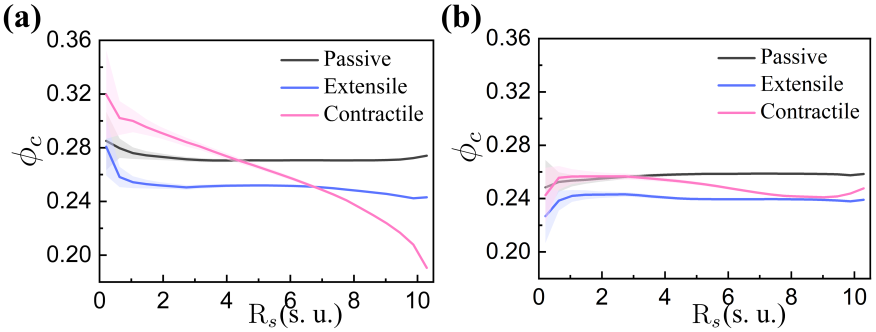


Fig. S3: *Chromatin density profiles for passive and motor cases without crosslinks and with chromatin–lamina linkages.* Chromatin density profiles, $\phi_c$, as a function of radial distance R$_s$ for passive, extensile, and contractile motor cases. (a) The system without crosslinks or chromatin-lamina linkages ($N_c = 0$, $N_L = 0$). (b) The system with chromatin-lamina linkages present but no crosslinks ($N_c = 0$, $N_L = 600$). All quantities are reported in simulation units (s. u.), where 1 s. u. of length = 1 $\mu m$ and 1 s. u. of time = 0.5 s.

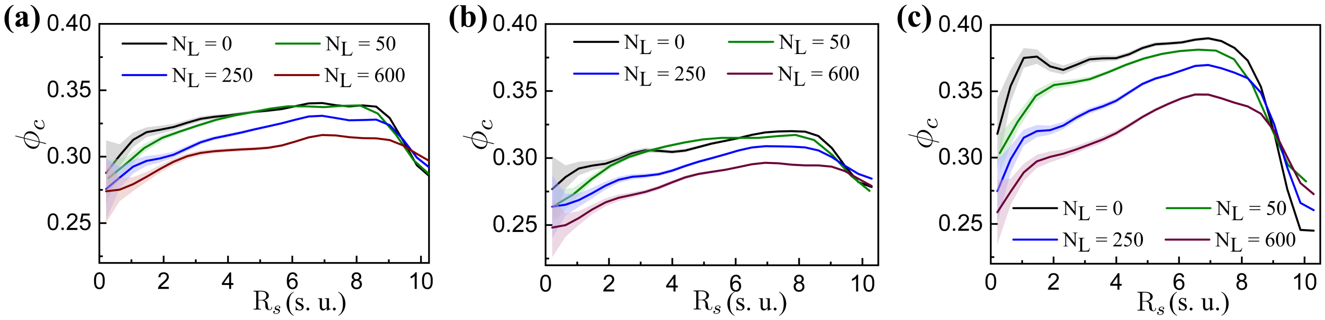


Fig. S4: *Chromatin distributions for passive, extensile, and contractile motors with varying chromatin–lamina linkages.* Chromatin density, $\phi_c$, as a function of R$_s$ for (a) passive, (b) extensile, and (c) contractile motors, shown for different chromatin-lamina linkage numbers ($N_L$) with a linear crosslink density profile. Results correspond to $N_c = 2000$ chromatin crosslinks. All quantities are reported in simulation units (s. u.), where 1 s. u. of length = 1 $\mu m$ and 1 s. u. of time = 0.5 s.

Next, we investigate how chromatin distribution is influenced by varying the number of chromatin–lamina linkages while keeping the crosslink profile linear and the total number of chromatin crosslinks fixed at $N_c = 2000$. Figure S4 shows the resulting chromatin density profiles under passive, extensile, and contractile motor conditions. Both the passive and extensile cases yield nearly uniform radial density profiles, with extensile motor activity promoting chromatin decompaction relative to the passive and contractile cases (Fig. S4(a, b)). In contrast, contractile motors lead to significant chromatin compaction, resulting in a pronounced increase in density near the nuclear periphery. When no chromatin–lamina linkages are present, or when their number is low, the chromatin distribution remains relatively uniform, with slightly elevated density throughout the nucleus compared to cases with a higher number of linkages (Fig. S4c). However, increasing the number of linkages markedly enhances the spatial differentiation of chromatin density, with lower density in the nuclear interior and accumulation at the periphery (Fig. S4c).

#### 2.3. Spatial distribution and dynamics of euchromatin and heterochromatin

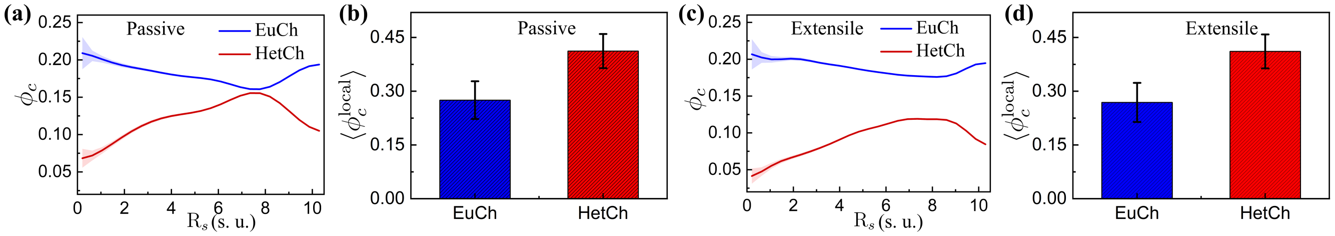


Fig. S5: *Euchromatin and heterochromatin density profiles for passive and extensile motor cases with a linear crosslink profile.* Density ($\phi_c$) of euchromatin (EuCh) and heterochromatin (HetCh) as a function of R$_s$, along with the average chromatin density ($\langle\phi_c\rangle$), for passive (a, b) and extensile motor (c, d) cases, for a linear crosslink density profile. The results correspond to $N_c = 2000$ chromatin crosslinks and $N_L = 600$ chromatin-lamina linkages. All quantities are reported in simulation units (s. u.), where 1 s. u. of length = 1 $\mu m$ and 1 s. u. of time = 0.5 s.

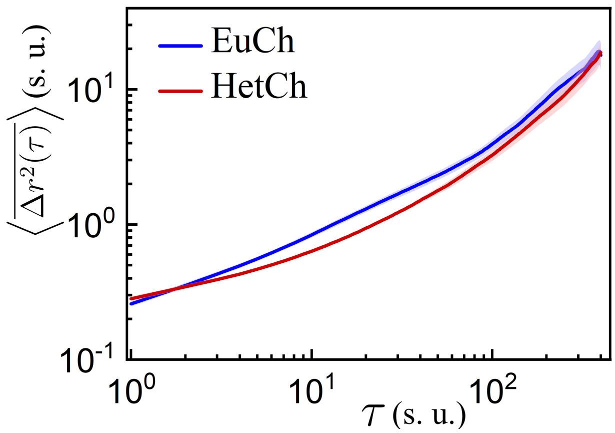


Fig. S6: *Dynamics of euchromatin and heterochromatin for contractile motors with a linear crosslink profile.* Mean square displacement $\left(\langle\overline{\Delta r^2(\tau)}\rangle\right)$ of euchromatin (EuCh) and heterochromatin (HetCh) for contractile motors with a linear crosslink density profile. The results correspond to $N_c = 2000$ chromatin crosslinks and $N_L = 600$ chromatin-lamina linkages. All quantities are reported in simulation units (s. u.), where 1 s. u. of length = 1 $\mu m$ and 1 s. u. of time = 0.5 s.

#### 2.4. Nuclear lamina shape fluctuations

To characterize shape fluctuations of the shell, a random slab passing through the center is selected, and the coordinates of the shell monomers within the slab are projected onto its plane. The spatial deviations of these monomers from the average shell radius are then analyzed using a fast Fourier transform (FFT), with $F_q$ denoting the Fourier component of the deviation at wavenumber $q$. Fig. S7 presents the power spectrum of shape fluctuations for passive, extensile, and contractile motors with a linear crosslink profile, and for different crosslink density profiles corresponding to passive, extensile, and contractile motor cases. Contractile motors exhibit large-amplitude shape fluctuations, indicating more pronounced deformation of the nuclear lamina compared to extensile and passive cases (Fig. S7a). However, for a given motor case, qualitatively similar trends are observed across different crosslink density profiles (Fig. S7(b-d)).

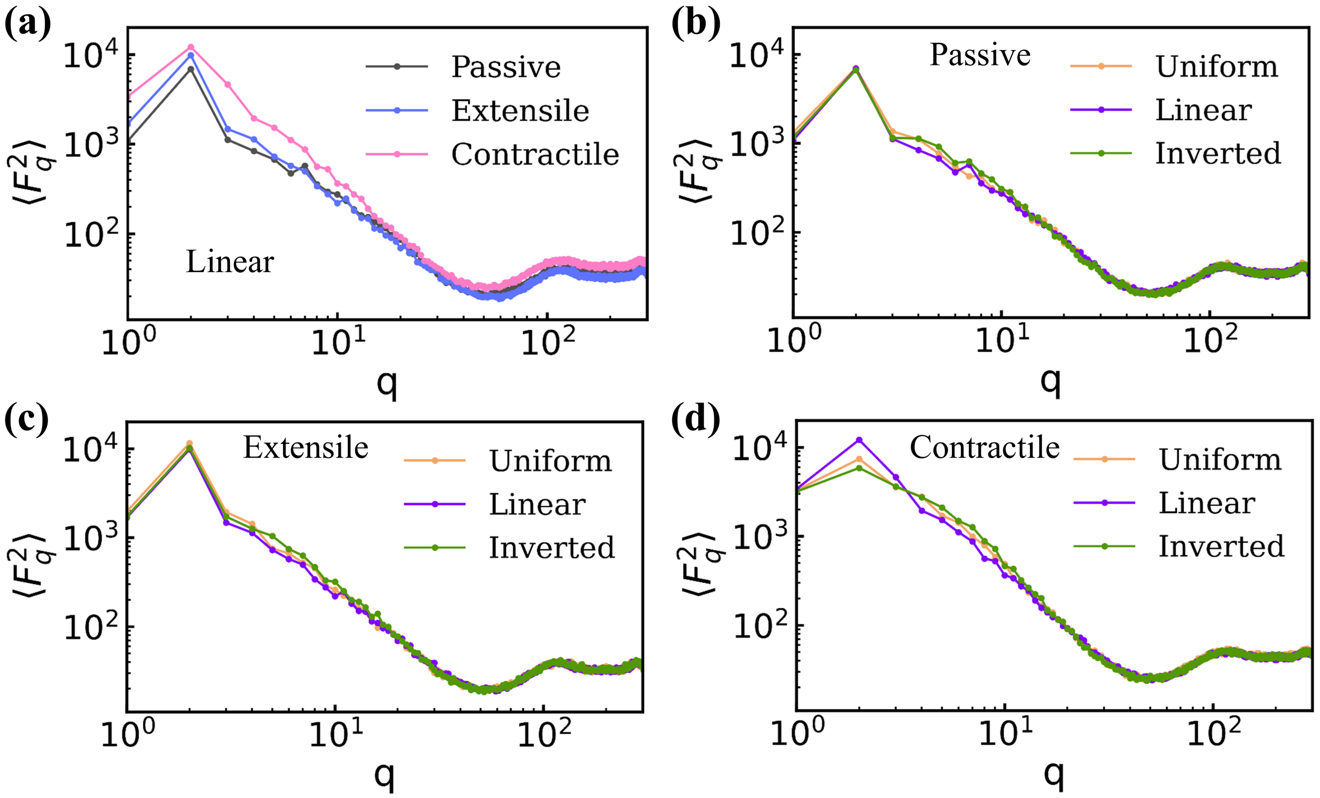


Fig. S7: *Nuclear shape fluctuations for different motor types and crosslink profiles.* Power spectrum of shell shape fluctuations for simulations with $N_c = 2000$ chromatin crosslinks and $N_L = 600$ chromatin-lamina linkages. (a) Different motor cases (passive, extensile, contractile) for a linear crosslink density profile. (b–d) Comparison of crosslink density profiles for (b) passive, (c) extensile, and (d) contractile motor cases.
